# Dandy Walker-like malformations in mutant mice demonstrate a role for PDGF-C/PDGFRα signalling in cerebellar development

**DOI:** 10.1101/2022.05.13.491787

**Authors:** Sara Gillnäs, Radiosa Gallini, Liqun He, Christer Betsholtz, Johanna Andrae

## Abstract

Formation of the mouse cerebellum is initiated in the embryo and continues for a few weeks after birth. Double mutant mice lacking platelet-derived growth factor-C and that are heterozygous for platelet-derived growth factor receptor alpha (*Pdgfc-/-; Pdgfra^GFP/+^*) develop cerebellar hypoplasia and malformation with loss of cerebellar lobes in the posterior vermis. This phenotype is similar to those observed in *Foxc1* mutant mice and the human syndrome Dandy Walker malformation. *Pdgfc-Pdgfra* mutant mice also display ependymal denudation in the 4^th^ ventricle and gene expression changes in cerebellar meninges, which coincide with the first visible signs of cerebellar malformation. Our observations suggest that PDGF-C/PDGFRα signalling is a critical component of the network of molecular and cellular interactions that take place between the developing meninges and neural tissues, and which are required to build a fully functioning cerebellum.

**Summary statement:** Mice lacking PDGF-C develop cerebellar hypoplasia and malformation. In addition, the ventricular zone close to the rhombic lip suffer from ependymal denudation.

## Introduction

Platelet-derived growth factors (PDGFs) play a number of important roles during vertebrate development and organogenesis (*Andrae et al., 2008*). The PDGF family consist of four different ligands (PDGF-A, -B, -C and -D) that signal through two tyrosine kinase receptors (PDGFR-α and -β). PDGF-C binds primarily to PDGFR-α (*Li et al., 2000*). We have previously shown that expression of PDGF-C (platelet-derived growth factor-C) is needed for proper formation of the cerebral meninges. Genetically modified mice lacking PDGF-C (*Pdgfc^-/-^; Pdgfra^GFP/+^*) develop thin and malformed meninges in combination with neuronal over-migration from the cerebral cortex into the leptomeningeal space and spinal cord defects (spina bifida) in newborn mice (*Andrae et al., 2016*).

The cerebellar development in mice starts already around E8-9 with the formation of the isthmic organizer. The process continues several weeks postnatally with the final positioning of neurons in the different cell layers and the foliation that generates the typical pattern of the cerebellum. Each step in the process is dependent on specific genes, and a multitude of cerebellar malformations has been identified both in mice and in humans (*Aldinger and Doherty, 2016; Haldipur and Millen, 2019*). Cerebellar malformations are commonly accompanied with malformations also elsewhere in the brain.

The human syndrome Dandy Walker malformation (DWM) affects the developing cerebellum and is characterized by cerebellar hypoplasia and upwards and dorsal rotation of the vermis (midline structure) in combination with enlargement of the fourth ventricle (*Parisi and Dobyns, 2003*). Dysplasia of the most posterior cerebellar lobes, referred to as vermian tail, can be observed in DWM using foetal MRI. The tail sign has been suggested to be specific for the DWM, but several questions remain unanswered regarding its clinical significance (*Bernardo et al., 2015; Teresa and P, 2018*). Although much is still to learn about genetic alterations that are causative of DWM, *Foxc1* mutant mice have been suggested as a model for DWM (*Aldinger and Doherty, 2016; Aldinger et al., 2009; Haldipur et al., 2017*) because they display similar cerebellar abnormalities, including defective cerebellar foliation, hypoplasia of the posterior vermis and defective layering of Purkinje cells (*Haldipur et al., 2017). Foxc1* encodes a transcription factor (Foxc1) that regulates expression of SDF-1 (*Cxcl12*) in meningeal cells (*Zarbalis et al., 2007*), which in turn bind to CXCR4 on granule cells in the external granular layer. Loss of SDF-1/CXCR4 interaction causes cerebellar malformations due to defective migration of cells in the external granular layer (EGL) cells and Purkinje cells (*Vilz et al., 2005)(Aldinger et al., 2009; Haldipur et al., 2017*).

Here we describe cerebellar malformation including hypoplasia and mis-location of the posterior vermis in *Pdgfc*^-/-^: *Pdgfra*^GFP/+^ mice. The phenotype is similar to that reported for *Foxc1* mutant mice, and we therefore suggest that also PDGF-C has a role in DWM. In addition, we show that denudation of the ependyma in the fourth ventricle is associated with the cerebellar phenotype in *Pdgfc^-/-^*: *Pdgfra^GFP/+^* mice.

## Results

We found cerebellar hypoplasia combined with a specific loss of the posterior inferior vermis in *Pdgfc^-/-^*: *Pdgfra^GFP/+^* mice. The phenotype was only observed in the combination of complete loss of *Pdgfc* and heterozygous loss of *Pdgfra*, suggesting that both Pdgfc and Pdgfa signaling via Pdgfra are important for cerebellar development. Abnormal cerebellar morphology was primarily seen the caudal lobes closest to the rhombic lip at the sagittal midline (Fig.1). All late embryos and newborn *Pdgfc^-/-^*: *Pdgfra^GFP/+^* mice were affected (Fig. 2). The severity of the malformation varied slightly between individuals, which was in line with some mice surviving a few days longer than others. *Pdgfc^-/-^*; *Pdgfra^GFP/+^* mice show several different developmental defects, but the exact cause of death is still not known (*Andrae et al., 2016*). Most mutant mice died at birth or had to be sacrificed within a few days postnatally. As we have reported earlier (*Andrae et al., 2016), Pdgfc^-/-^; Pdgfra^GFP/+^* individuals were readily spotted in newborn litters of mixed genotypes by the presence of a visible red stripe in the skin above the caudal spinal cord; this stripe is associated with an underlying spina bifida.

**Fig. 1.**
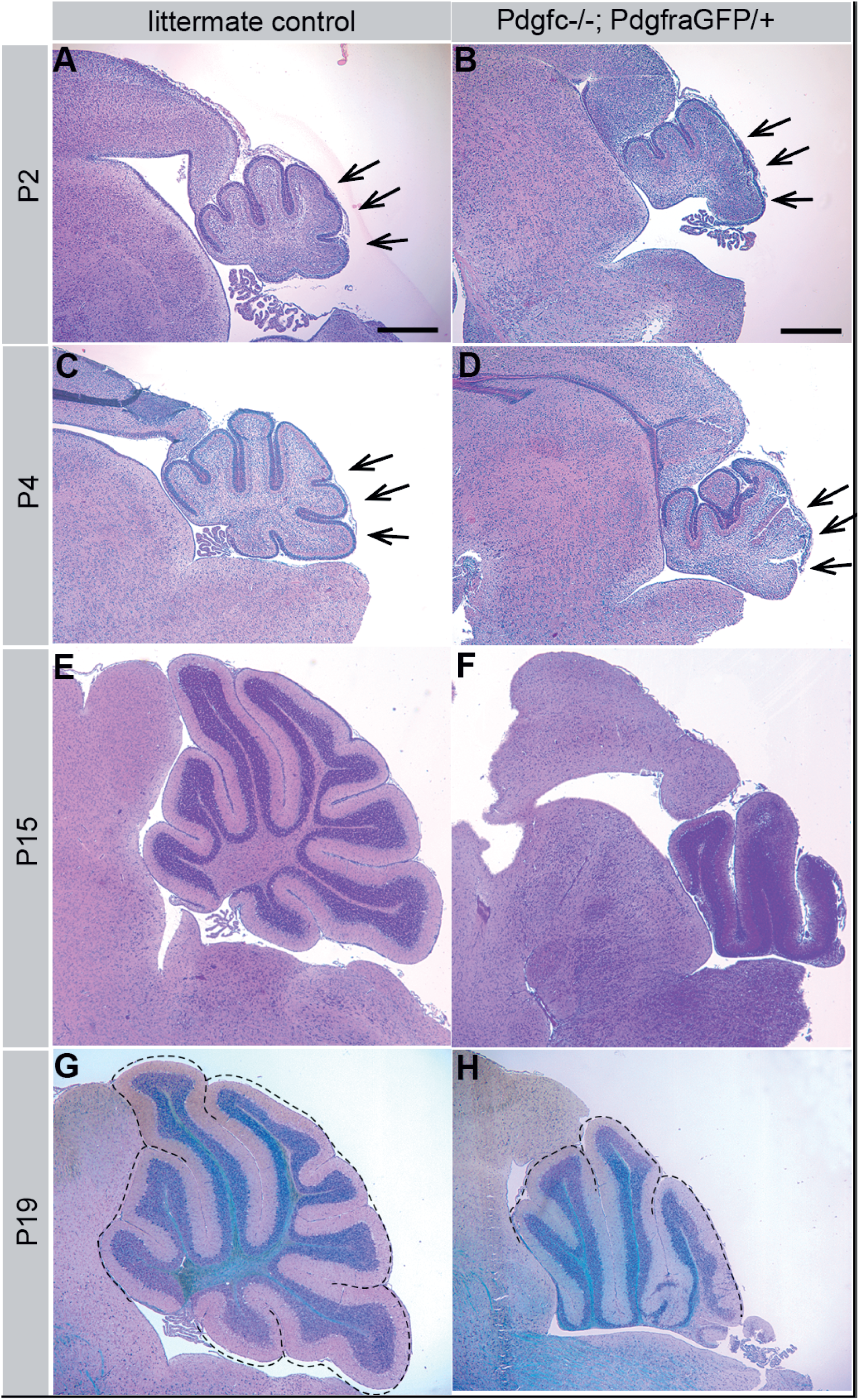
Cerebellar malformation in postnatal *Pdgfc^-/-^; Pdgfra^GFP/+^* mice. Hematoxylin-eosin staining of mid-sagittal sections from wild type cerebellum at (A) P2, (C) P4, (E) P15 and (G) P19. Hypoplasia and loss of caudal lobes in cerebellum of *Pdgfc^-/-^; Pdgfra^GFP/+^* mice at (B) P2, (D) P4, (F) P15 and (H) P19. Dotted line in G and H marks the cardinal lobes. Arrows indicate area where lobes were missing/malformed. Several P2 and P4 brains were analysed with similar morphology. Brains from P15 and P19 represent individual samples.

**Fig. 2.**
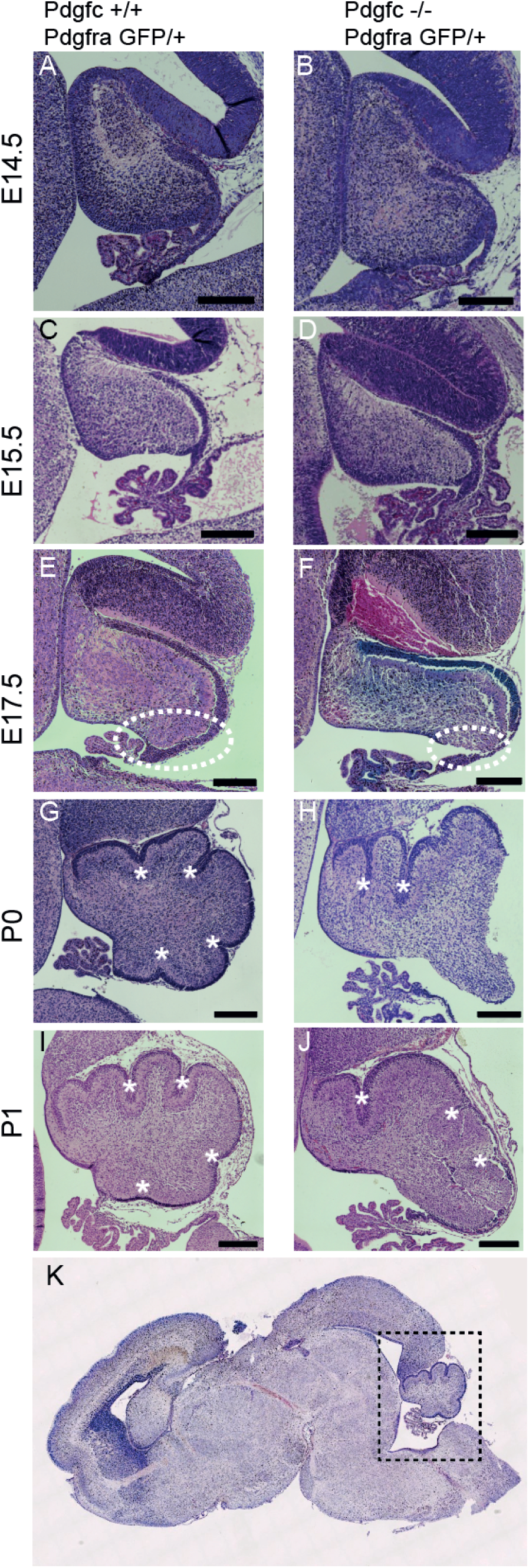
Embryonic development of cerebellum in *Pdgfc^-/-^; Pdgfra^GFP/+^* mice. Hematoxylin-eosin staining of mid-sagittal sections of cerebellum. No morphological differences between mutant and control mice at E14.5 (A, B) or E15.5 (C, D). At E17.5 the rhombic lip area (encircled) has expanded in control mice (E), but not in mutant mice (F). From birth, the primary fissures (asterisk) and the cardinal lobes have formed in control mice (G, I). In mutant mice (H, J), only the rostral lobes are clearly visible. (K) Dotted lines indicate a mid-sagittal view of the cerebellum. All sections are representative of >3 analysed brains. Scalebar 200 *μ*m.

### Cerebellar hypoplasia and dis-organisation of neurons

The first signs of malformations in the cerebellum of *Pdgfc^-/-^; Pdgfra^GFP/+^* mice were observed at E15-16, in the area close to the rhombic lip (Fig. S1A, B). In control mice, the cerebellum expanded into a rounded appearance and as a consequence, the choroid plexus and the rhombic lip rotated ventrally into the fourth ventricle (Fig. 2E). The *Pdgfc^-/-^*; *Pdgfra^GFP/+^* cerebellum instead displayed a flat and elongated shape with the rhombic lip positioned dorsally (Fig. 2F). At birth, the four primary fissures and the five cardinal lobes had started to form in control mice and were clearly visible at the sagittal midline (Fig. 2G, I). In mutant mice, the two rostral cardinal lobes appeared normal and a central lobe was present but had an abnormal shape. The caudal lobes were, however, essentially missing, which changed the position of the rhombic lip to a more dorsal position, away from the fourth ventricle (asterisk in Fig. S1F). A similar phenotype has been reported in *Foxc1* mutant mice, where it was suggested to model the DWM syndrome (*Haldipur et al., 2017*).

A limited number of *Pdgfc*^-/-^; *Pdgfra^GFP/+^* mice displayed a somewhat less severe phenotype, which gave us the possibility to analyse the cellular organisation also in postnatal brains. The overall organization of granular-, molecular- and Purkinje cell layer and the white matter were largely intact in the rostral lobes (Fig. 1). At more posterior location in the midline vermis, the neuronal layers appeared progressively disorganized, involving all cell layers. Ectopic clusters of granular cells, irregular multicellular layers of Purkinje cells, and an uneven molecular layer were observed (Fig.3). The position of Purkinje cells in *Pdgfc^-/-^; Pdgfra^GFP/+^* mice was abnormal already at birth, where the multicellular layer of calbindin-positive Purkinje cells was uneven and did not follow the shape of the developing lobes (Fig. 3A, D). At P15, abnormal clusters of Purkinje cells were still observed in some areas, whereas other areas had a more normal appearance with Purkinje cells positioned one-by-one (Fig. 3C, F). In the caudal cerebellum, which displayed the highest degree of cellular disorganisation, the pattern of cell nuclei was completely disrupted (Fig. 3L). Bergmann glia, lacked their typical perpendicular orientation in mutants and were regionally missing and glial endfeet were absent (Fig. 3L’).

**Fig. 3.**
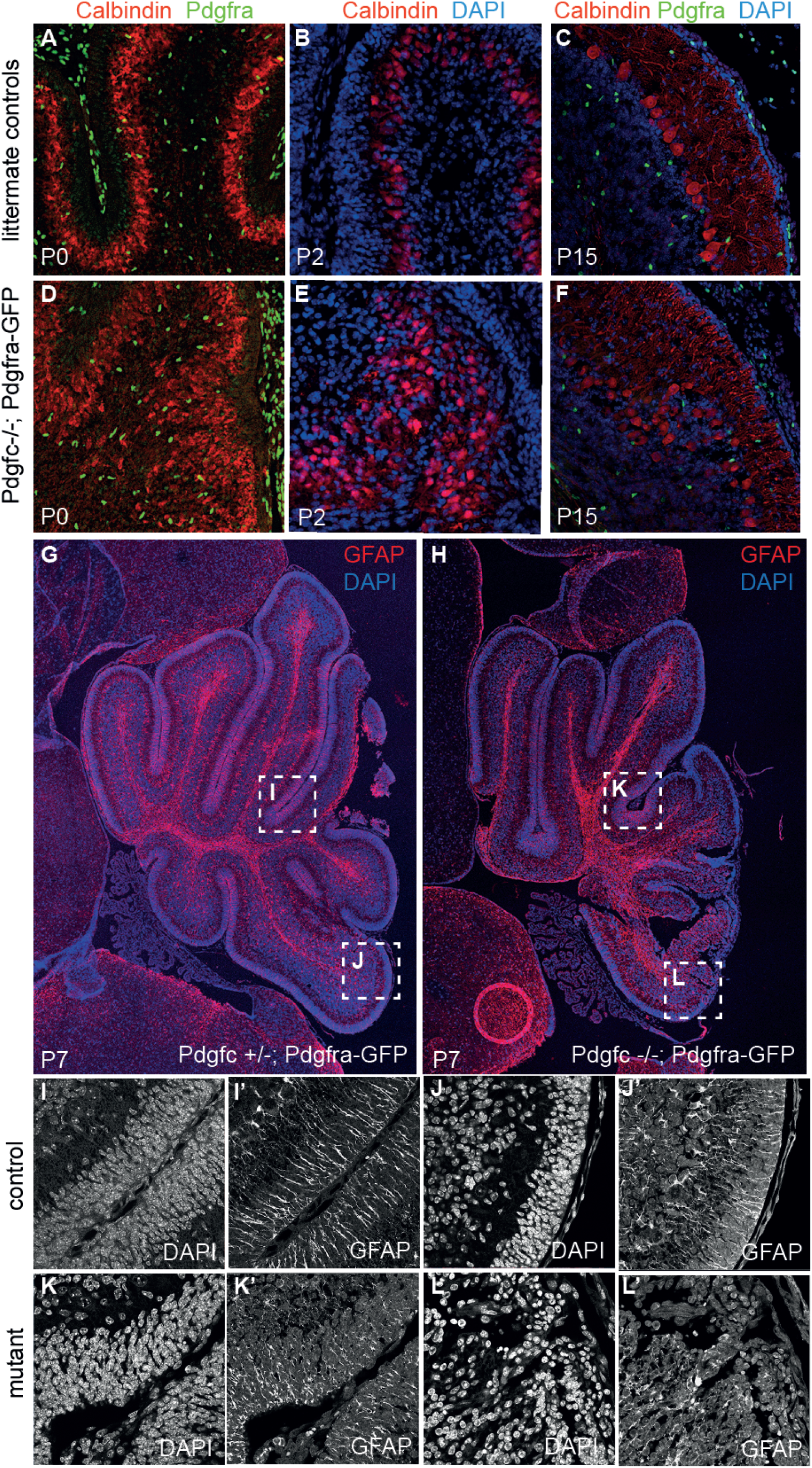
Irregular pattern of Purkinje cells and Bergman glia. Immunofluorescent staining of cerebellum from postnatal mice. (A-C) Calbindin marks how Purkinje cells in control mice were evenly distributed at P0 and acquired a single cell layer at P15. (D-F) Purkinje cells in mutant mice were unevenly distributed with partly ectopic expression. (G) P7 control mouse. (H) P7 *Pdgfc^-/-^; Pdgfra^GFP/+^* mouse. (I-J) Evenly distributed cell nuclei and perpendicular Bergman glia in both rostral and caudal area of control mice. (K) Evenly distributed cell nuclei, but partly crocked Bergman glia in rostral area of P7 mutant mice. (L) Disorganized pattern of cell nuclei and no perpendicular Bergman glia. Several P0 and P2 brains were analysed. P7 is an individual sample and P15 is a representative of two.

### Ependymal disruption close to the choroid plexus in the 4^th^ ventricle

As a consequence of the mis-location and dorsal positioning of the rhombic lip area the ventricular lining from the colliculus to the choroid plexus in the 4^th^ ventricle was elongated in *Pdgfc^-/-^; Pdgfra^GFP/+^* mice compared to the control mice (yellow dotted line, Fig. 4A, B). In addition, the cells lining the ventricular zone (VZ) closest to the rhombic lip (RL) in the mutant mice did not organize into a continuous layer of polarised ependymal cells (Fig. 4C, D). This lost ependymal layer integrity was visible by hematoxylin-eosin staining already at embryonal day 17.5 (E17.5) (encircled area in Fig. 2F). The mutant ependymal cells in this area had lost their epithelial-like morphology and was completely lacking apical podocalyxin (*Podxl*) expression (Fig. 4C’, D’). Loss of podocalyxin expression was largely restricted to the sagittal midline and was not seen in lateral sections (data not shown). To confirm the change in identity of the cells lining the ventricular zone, we used ependymal markers including microtubular protein βIV-tubulin (expressed in the cilia of the ependymal cells) and the cytosolic calcium-binding protein S100B (Fig. 5). Neither βIV-tubulin nor S100B were observed at the 4^th^ ventricle midline in mutants, i.e. the region also lacking *Podxl* expression (Fig. 5B, D), which further confirms the loss of ependymal layer integrity in this region. With the observed disrupted ependyma in *Pdgfc^-/-^; Pdgfra^GFP/+^mice*, the brain parenchyma close to the rhombic lip would suffer direct exposure to cerebrospinal fluid present in the 4^th^ ventricle. Disruption of the ependymal cell layer is also referred to as denudation (*Sival et al., 2011*). Previous studies have observed how denudation activates a repair mechanism by accumulation of astroglial processes (*Sarnat, 1995*) and thereby upregulate expression of glial fibrillary acidic protein (GFAP). In *Pdgfc^-/-^; Pdgfra^GFP/+^* mice, GFAP was indeed upregulated and covered the area with disrupted ependymal cells (Fig. 5F), suggesting activated repair involving GFAP-positive astrocyte processes. We also analyzed the expression of additional ependymal markers and junction proteins in the mutant mice (Fig. S2). Changes in expression of the neuroepithelial junction protein n-cadherin (Cdh2) and the gap-junction protein connexin 43 (Cx43) were observed from E17.5 in cells lacking podocalyxin (Fig. S2D). Earlier, at E15.5, we did not observe abnormal ependymal denudation and marker expression in this area.

**Fig. 4.**
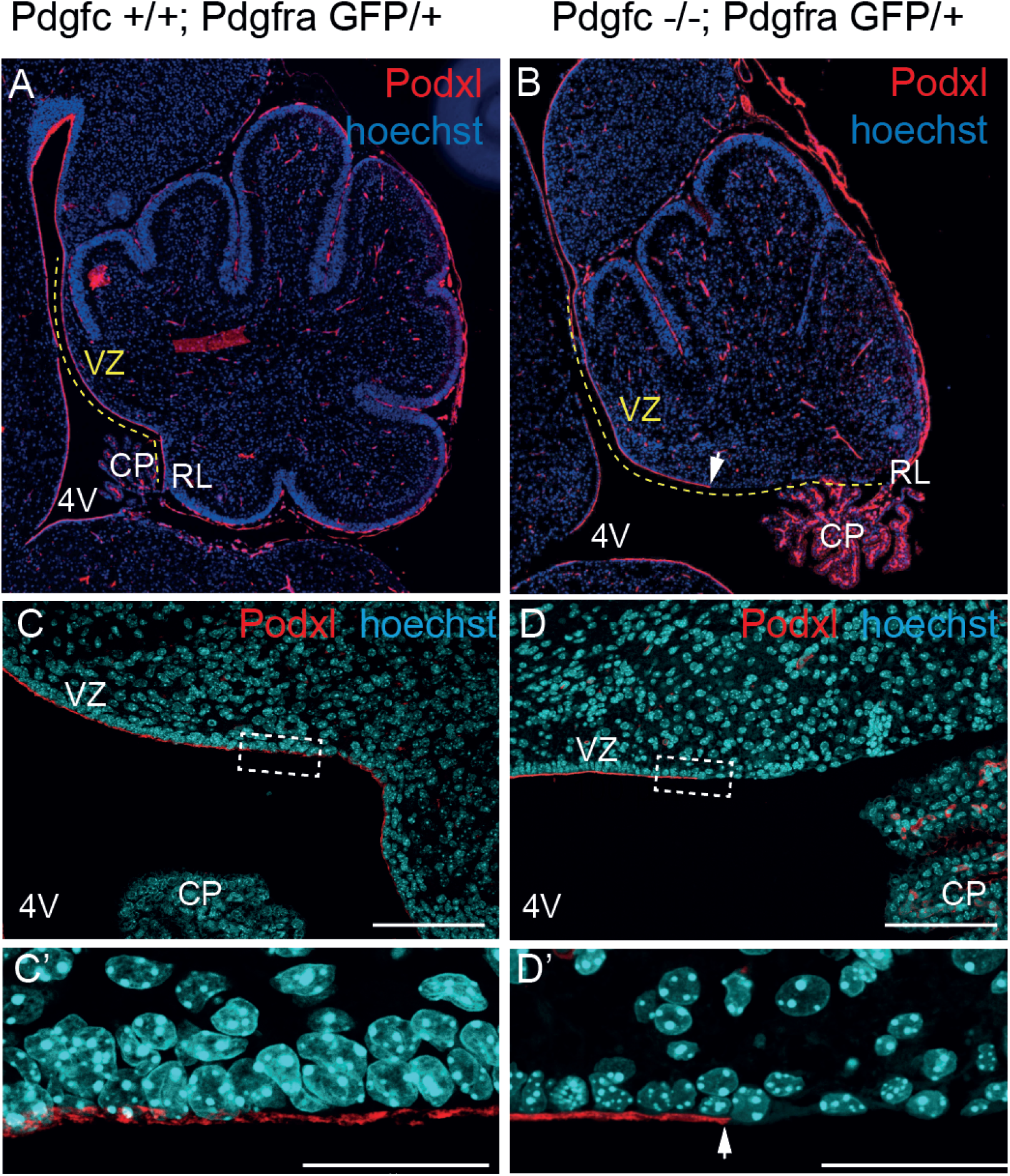
Lost integrity of ependymal cells. Immunofluorescent staining of podocalyxin at P1. (A) Podocalyxin was expressed along the ventricular zone of the control cerebellum. (B) In the mutant cerebellum, podocalyxin was not expressed in the ventricular zone closest to the rhombic lip. The ventricular zone was also extended as the rhombic lip and choroid plexus rotated dorsally. (C) Cuboidal ependymal cells in control mice line up side-by-side with apical expression of podocalyxin. (D) In mutant mice, the regular pattern of ependymal cells was lost, and the podocalyxin expression ended distinctly before reaching the rhombic lip. VZ-ventricular zone, CP-choroid plexus, 4V-fourth ventricle, RL-rhombic lip. Arrows indicate where podocalyxin expression was lost. Scalebar (C, D) 100 *μ*m, (C’, D’) 30 *μ*m. Images are representatives of stainings from > 3 mice.

**Fig. 5.**
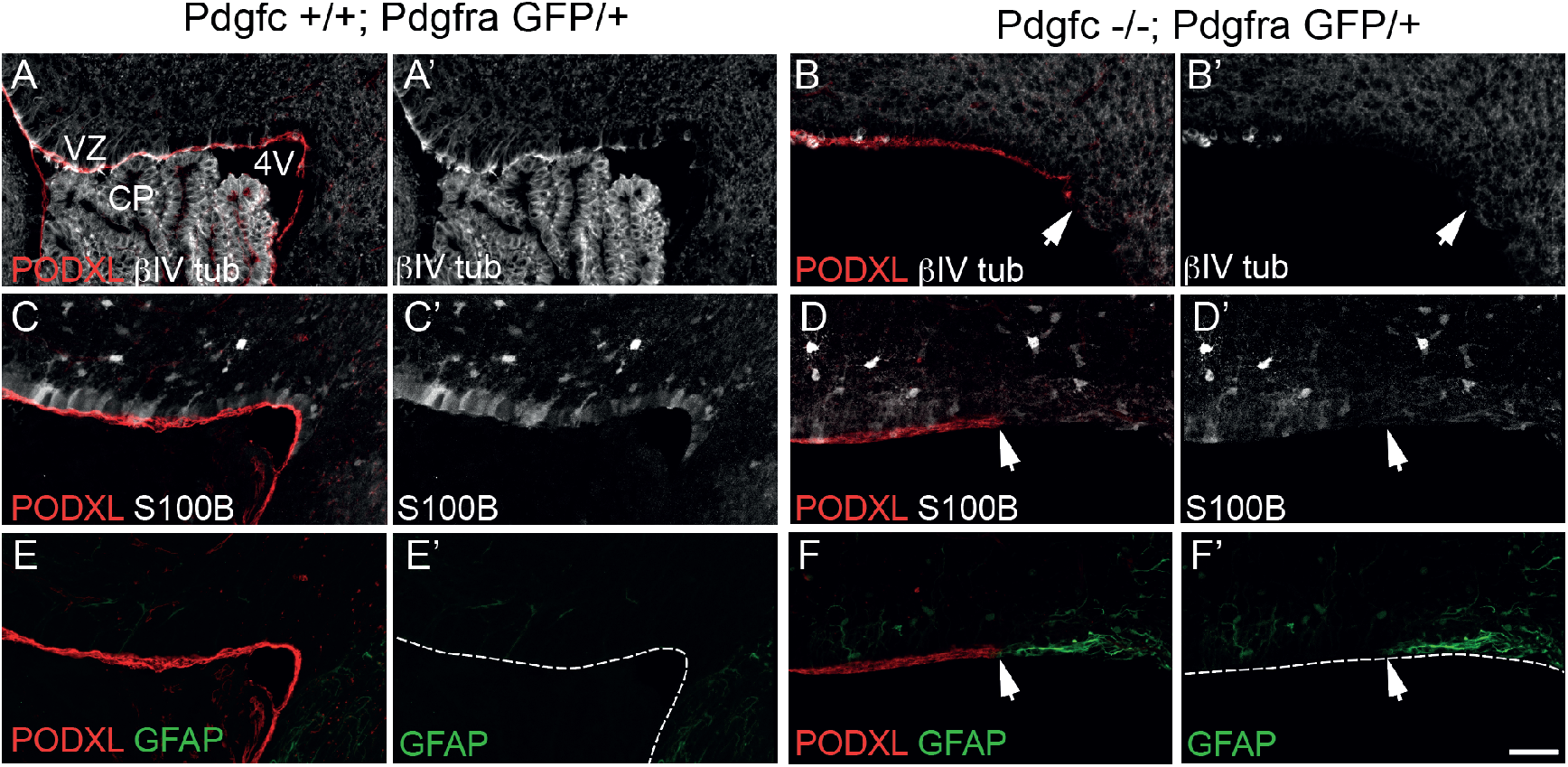
Upregulation of GFAP in areas of denudation. Immunofluorescence stainings of podocalyxin, βIV-tubulin, S100B and GFAP in ventricular zone at P0. (A, B) Expression of βIV-tubulin was lost in mutant cells lacking podocalyxin expression. (C, D) Expression of S100B was reduced in mutant cells lacking podocalyxin expression. (E, F) Lack of podocalyxin was replaced by GFAP in mutant mice. VZ-ventricular zone, CP-choroid plexus, 4V-fourth ventricle. Dotted line indicates border of the ventricular zone. Arrows indicate where podocalyxin expression was lost. Scalebar 50 *μ*m. Images are representatives of stainings from > 3 mice.

### *Pdgfc* expression in the developing cerebellum

As shown above, PDGF-C is necessary for proper development of the cerebellum. To better understand the process, we investigated the time and anatomical location of the onset of *Pdgfc* expression in the cerebellum. We took advantage of the *Pdgfc* knockout mice, in which introduction of a *lacZ* gene was used to generate the *Pdgfc* null allele (Ding et al., 2004). X-gal staining in the hindbrain of mutant mice (*Pdgfc^-/-^; Pdgfra^GFP/+^*) was compared to heterozygous control mice (*Pdgfc^+/-^; Pdgfra^GFP/+^*) from E9.5 until birth. *Pdgfc-lacZ* was expressed in the notochord and floor plate close to the hindbrain as early as E9.5 (data not shown). Within the cerebellar tissue, the first expression was visualized in the caudal part of the cerebellar primordium at E12.5 (Fig. A-C). Interestingly, at this stage the only putative target cells of PDGF-C (i.e. cells expressing *Pdgfra*) were positioned in the overlying mesenchyme (Fig. 6G, H). In mutant embryos, X-gal staining was stronger than in control embryos. Likely, this was due to the presence of double copies of *lacZ* in the mutant mice (Fig. 6D-F). At E16.5, when cerebella in *Pdgfc^-/-^; Pdgfra^GFP/+^* embryos could be morphologically distinguishable from control embryos, strong lacZ expression was seen in granule cells of the transient EGL. At this stage, EGL cells are known to migrate anteriorly from the rhombic lip to cover the newly fused cerebellar anlagen (Fig. S1). Mutant embryos displayed a more widespread expression pattern suggesting ectopic migration of *Pdgfc*-expressing cells or, alternatively, the compensatory upregulation of *Pdgfc* by other cell types (Fig. S1B, D, F)

**Fig. 6.**
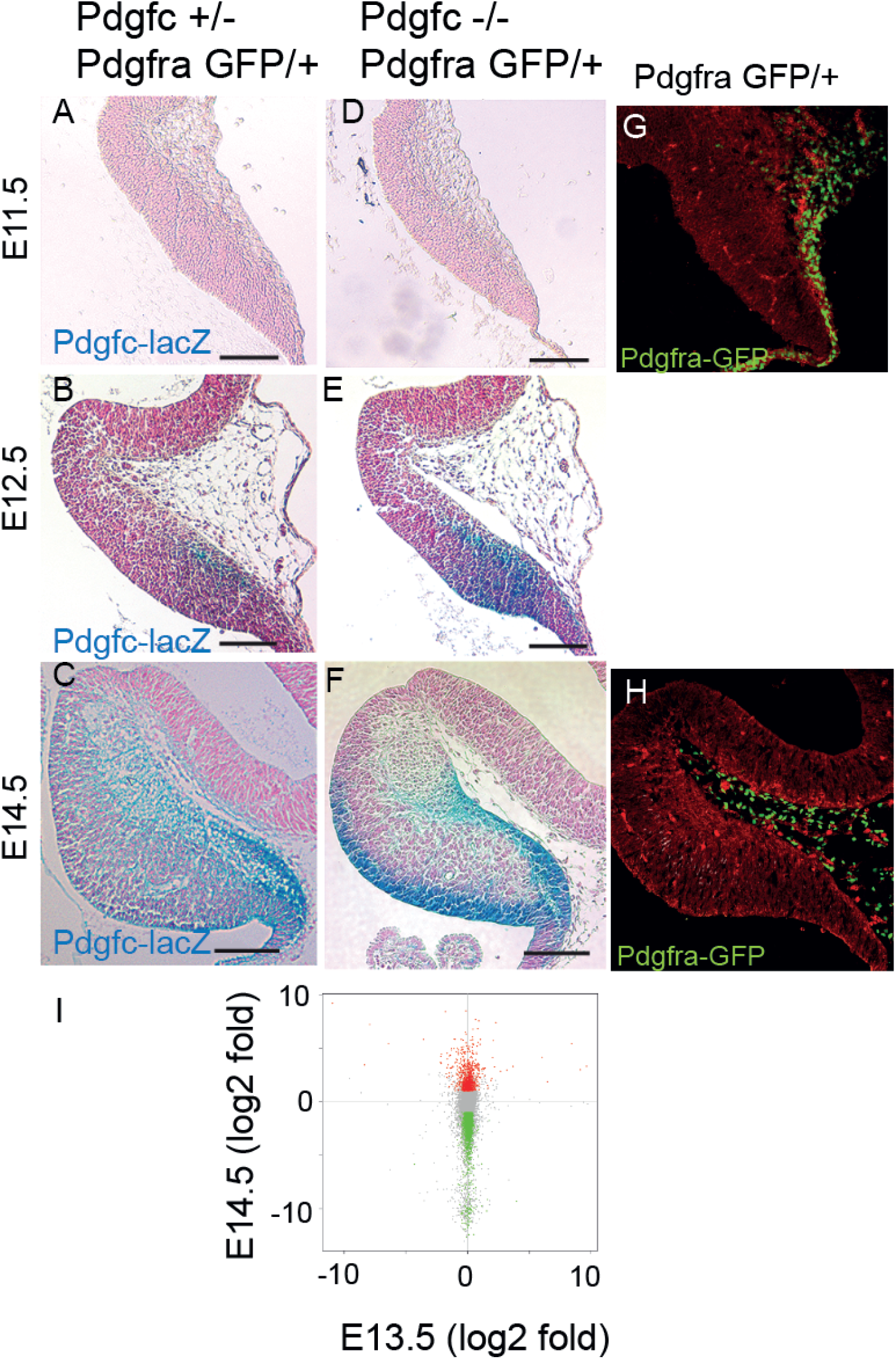
Embryonic expression of *Pdgfc* and gene expression changes in mutant mice. (A-H) X-gal staining reflecting *Pdgfc* promoter activity and *Pdgfra*-GFP reporter expression in cerebellar primordium at E11.5, E12.5 and E14.5. Each section is a representative of >3 mice. (A-C) *Pdgfc* promoter activity in control mice was first identified at E12.5. At E14.5, strong expression was seen in the rhombic lip area (D-F) X-gal staining was stronger in mutant mice. (G, H) *Pdgfra*-GFP reporter was observed in the overlying meninges, but not within the cerebellar tissue itself. (I) Differentially expressed genes identified by microarray analyses of cerebellar tissue in mutant and control mice at E13.5 and E14.5. No variation was seen at E13.5 (x-axis), whereas E14.5 (y-axis) display several significantly regulated genes. Statistically significant regulated genes (p<0.05 and fold change >2) are shown in red (up-regulated) and green (down-regulated). Non-significant genes are in grey.

### Altered gene expression in the absence of PDGF-C

We wanted to identify changes in gene expression that were associated to the loss of PDGF-C in the developing cerebellum. Knowing that the *Pdgfc* expression in the cerebellar area was initiated around E12.5 (Fig. 6C) and morphological abnormalities were observed around E15.5 (Fig. S1A), we hypothesized that secondary gene expression changes resulting from the loss of PDGF-C signalling might occur between E12,5 and E15.5. Microarray gene expression analyses were therefore performed on RNA from cerebellar tissue (liberated from meninges) at E13.5 and E14.5. At E13.5, gene expression was almost identical in *Pdgfc^-/-^; Pdgfra^GFP/+^* mutants and control pups. The limited number of differently regulated genes observed could be explained by the sex distribution in the litter. However, one day later (at E14.5), a significantly higher number of differentially expressed genes were identified in the *Pdgfc^-/-^; Pdgfra^GFP/+^* cerebellum compared to the controls (fold change >2, p<0.05) (Fig. 6I). Taken together, this fit well into the time frame with loss of *Pdgfc* expression at E12.5, transcriptional changes and altered gene expression around E14.5 and morphological changes around E15.5.

The large number of genes down-regulated at E14.5 indicated that PDGF-C is an important player in the developing cerebellum. Most of the GO terms were related to development and cellular processes, e.g. *G-protein coupled receptor signalling pathway, Sensory perception* and *Nervous system process* (supplementary Table S1). The top down-regulated genes at E14.5 were manually compared to scRNA seq data from the adult mouse brain (Zeisel et al., 2018). However, most of the genes indicated expression in different types of cells and among the down-regulated genes were also a large number of olfactory- and vomeronasal receptors, which we could not connect to specific cells. A few relevant genes were anyway identified; e.g. the ependyma-specific genes *Mak* and *Bbox1* were totally missing in mutant mice, the Purkinje cell marker *Calb1* was >5 fold down-regulated and meningeal markers as *Cxcl12* and *Crym* (DeSisto et al., 2020) were down-regulated -3.24 fold and -2.41 fold, respectively.

A third microarray analysis was performed at P0, this time on both cerebellar tissue and the overlying meninges from mutant and control mice. Very few significant gene expression changes were observed in the cerebellar tissue, but in the mutant meninges, there were clear gene expression changes. The differentially down-regulated genes were highly associated to *Vasculature development* and similar GO terms (supplementary Table S2). Among the differentially down-regulated genes in the meninges of *Pdgfc^-/-^; Pdgfra^GFP/+^* mice were e.g. *Col15a1* (−3.2 fold), *Fmod* (−2.75 fold), *Vnn1* (−4.18 fold) and *Slc22a2* (−3.19 fold). These genes reflect different meningeal fibroblasts (Del Gaudio, *manuscript in preparation*).

## Discussion

In this study we demonstrate that PDGF-C plays an important role during the development of the mouse cerebellum. Loss of the *Pdgfc* gene, and thereby reduced activation of the PDGFRα, affected the cerebellar formation already during the embryonal stage. Formation of the cerebellum is complex and in the *Pdgfc^-/-^; Pdgfra^GFP/+^* mice several distinct defects were identified; e.g. hypoplasia of the posterior vermis associated to a relative change in the position of the rhombic lip, lost cellular identity of ependymal cells in the 4^th^ ventricle, ectopic migration of EGL cells, and uneven and multicellular layers of Purkinje cells. As a result, postnatal mice presented cerebella with defect foliation and a complete lack of the most caudal lobes of the cerebellum.

Many processes occur from the start of *Pdgfc* expression (E12.5) in the developing embryonic cerebellum to the fully mature cerebellum postnatally. One interesting aspect is that whereas *Pdgfc* was expressed within the cerebellar tissue, we could only detect possible responding *Pdgfra* positive cells in the meninges and in the surrounding mesenchyme. The most distinct *Pdgfc* expression was in the EGL-cells. This pattern coheres with scRNAseq data from the developing human cerebellum, where *PDGFC* is expressed in EGL cells and *PDGFRA* in oligodendrocyte progenitor cells, meninges and choroid plexus/ependyma (*Aldinger et al., 2021*).

Interactions between meningeal cells and cells in the developing brain are necessary for proper formation of the CNS. The importance of signalling between EGL cells and meningeal cells is well-known for proper foliation of the cerebellum (*Sievers et al., 1981*). *Foxc1* is only expressed in the meninges and still has such an important impact on formation both on the cerebellum (*Aldinger et al., 2009*) and on the cerebral cortex (*Siegenthaler et al., 2009*).

The cerebellar phenotype of the *Pdgfc^-/-^; Pdgfra^GFP/+^* mice showed great similarity to *Foxc1* mutant mice (*Haldipur et al., 2014; Haldipur et al., 2017*). FOXC1 loss in mice contributes to the formation of typical anomalies used to diagnose the human cerebellar malformation syndrome Dandy-Walker malformation; hypoplasia of the cerebellar vermis, upwards rotation of the vermis away from the brainstem and an enlargement of the 4^th^ ventricle (*Haldipur and Millen, 2019*). *Foxc1* is expressed in the meninges, where reduced levels of FOX1 leads to reduced levels of SDF-1 (Cxcl12) (*Zarbalis et al., 2007*). The levels of SDF-1 in the meninges were reduced also in newborn *Pdgfc^-/-^; Pdgfra^GFP/+^* mice. This further strengthen the hypothesis of a common signalling pathway responsible for the cerebellar phenotypes in *Pdgfc^-/-^; Pdgfra^GFP/+^* mice and in *Foxc1* mutant mice. In addition, a genetic interaction between *Foxc1* and *Pdgfra* has been indicated in a previous study on zebrafish, where *Pdgfra* was suggested to act downstream of *Foxc1 (French et al., 2014*).

The cellular interface between the fourth ventricle and brain parenchyma in the cerebellum was disrupted in *Pdgfc^-/-^; Pdgfra^GFP/+^* mice. The lack of intact ependymal cells was confirmed as the polarized *Podxl* expression was replaced by upregulated *Gfap* expression in the ventricular zone close to the rhombic lip. Ependymal disruptions (also called ependymal denudation) can be explained e.g. by mechanical forces and by increased pressure in the ventricular system. Ependymal cells are generated from radial glial cells (*Spassky et al., 2005*). The morphology and position of Bergman glia were disrupted in other areas of the *Pdgfc^-/-^; Pdgfra^GFP/+^* mice, hence migration defects could be one possible explanation. Further, it has been suggested that alterations in *Cdh2* (N-cadherin) expression proceeds ependymal denudation (*Oliver et al., 2013*). N-cadherin is needed to keep the ependymal layer intact and to maintain communication between the ependymal cells, and N-cadherin expression was altered in the *Pdgfc^-/-^; Pdgfra^GFP/+^* mice. Denudation of the ependyma in *Pdgfc^-/-^; Pdgfra^GFP/+^* mice was consistently and exclusively observed in the normally acute angle of the ventricular zone, shortly after the first signs of vermis hypoplasia and mis-location of the rhombic lip. Hence, another possible explanation is that mechanical stretching and tearing, enforced by the rotation of the vermis, contributed to the disrupted ependyma.

We were able to visualize early gene expression changes in the developing cerebellum of E14.5 embryos. A drawback with microarray analysis of a whole tissue is obviously that gene expression changes in individual cell types are easily masked and hidden, and it was difficult to pin-point relevant down-regulated genes in the bulk-RNA study. A large number of the differently regulated genes were olfactory- and vomeronasal receptors. Expression of olfactory receptors out of the olfactory tissues (e.g. in the developing mouse cortex) have been detected in various gene expression analyses, (*Feldmesser et al., 2006; Otaki et al., 2004*).

In summary, we present a cerebellar phenotype in the *Pdgfc^-/-^; Pdgfra^GFP/+^* mice, which show great similarities to the *Foxc1* mutant mice. We suggest that PDGF-C has a role in the complex network required to build a proper cerebellum and in the formation of the Dandy-Walker syndrome.

## Material and Methods

### Mouse strains and ethical permissions

All experiments were in accordance to Swedish animal welfare legislation. Ethical permits were approved by the regional research animal ethics committee in Stockholm (N24/06, N72/07, N33/10, N15/12) and in Uppsala (C225-12, C115/15, 5.8.18-03029-2020). Mouse strains used were: *Pdgfc* knockout mice (Pdgfc^tm1Nagy^, (Ding et al., 2004) where the lacZ gene replaces *Pdgfc;* and *Pdgfra^GFP/+^* (Pdgfra^tm11(EGFP)Sor^, (Hamilton et al., 2003) where expression of nuclear green fluorescent protein (GFP) replaces *Pdgfra*. The two strains were crossed to generate double transgenic mice *Pdgfc^-/-^; Pdgfra^GFP/+^*. Littermate *Pdgfc^+/+^; Pdgfra^GFP/+^* mice were used as controls, if not indicated differently. The majority of the mice were on a pure genetic background (C57BL/6J). Mice on a mixed genetic background (C57BL/6J;129S1) were also analysed and displayed an identical phenotype. Mutant *Pdgfc^-/-^; Pdgfra^GFP/+^* mice were easily recognized with a hemorrhagic stripe over the lower spine (Andrae et al., 2016). Genotyping for *Pdgfc* was carried out with PCR and expression of *Pdgfra*-GFP was confirmed using a fluorescence lamp. Both female and male mice were included in the study.

### Tissue processing

Embryos and newborn mice were sacrificed by decapitation, brains were dissected out, washed in PBS and fixed overnight in 4% paraformaldehyde (PFA). Mice older than one week were perfused through the heart with Hanks’ balanced salt solution (HBSS) and 4% PFA before dissection, and postfixation overnight. Fixed brains were either; dehydrated, embedded in paraffin and sectioned at 7 μm; or vibratome sectioned at 75 μm.

### Immunofluorescence staining (IF), histology and x-gal staining

Primary antibodies at the following concentrations were used: goat-anti-mouse Podxl (1:200, R&D Systems, AF1556); rabbit-anti-GFAP (1:100, Invitrogen, 13-0300); rabbit-anti-GFAP (1:200, Dako, Z0334); rabbit-anti-S100B (1:100, Dako, Z0311); rabbit-anti-βIV tubulin (1:500, Abcam, ab179509); rat-anti-Cdh2 (1:50, Developmental Studies Hybridoma Bank, MNCD2-C); mouse-anti-Cx43 (1:50, Santa Cruz Biotechnology, sc-271837), rabbit-anti-Calbindin (1:200, Sigma, D2724). Secondary antibodies were produced in donkey and conjugated to Alexa488, Cy3, Alexa 568 or A647 (Fischer Scientific). Additional antibody information in table S3. All stainings were performed together with negative controls, where the primary antibodies were excluded. All stainings were performed on tissue from at least three different animals.

Paraffin sections were deparaffinized in xylene and graded series of EtOH. Antigen retrieval in Target Retrieval Solution pH 9 (S2367, Dako, Glostrup, Denmark) was used for Podxl, Cdh2 and Cx43. Sections were washed in PBS, blocked in 1% bovine serum albumin (BSA), 0.5% Triton X-100 (T8787, Sigma), 2.5% normal donkey serum in PBS for a minimum of 30 min at room temperature (RT). Incubation with primary antibody was in 0.5% BSA, 0.25% Triton X-100 (T8787, Sigma) in PBS overnight at +4°C. Sections were washed in PBS, incubated with secondary antibodies for a minimum of 1h in RT, washed in PBS and stained with Hoechst (1:10 000 in PBS). Sections were mounted with ProLongGold antifade mountant.

Free-floating vibratome section were stained as above, but slowly shaking and with longer incubation times to allow for proper penetration into the tissue. Blocking was overnight, primary antibodies were incubated for 48h and secondary antibodies were incubated overnight. Sections were washed in PBS with 0.1% Tween 20 (P1379, Sigma).

X-gal staining was performed on *Pdgfc^-/-^; Pdgfra^GFP/+^* and *Pdgfc^+/-^; Pdgfra^GFP/+^* control mice. The method for x-gal-staining has been described before (*Andrae et al., 2014*). Up to E12.5, embryos were stained as whole mounts. From E13.5, brains were freely dissected before staining. X-gal-stained tissues were embedded in paraffin, sectioned at 7 *μ*m and counterstained with Nuclear Fast Red (N3020, Sigma Aldrich).

Hematoxylin and Eosin staining were performed on paraffin sections. Briefly, sections were deparaffinized, rehydrated and immersed in Mayer’s hematoxylin (HTX) (01820, HistoLab, Sweden) for 2min. Blueing was performed in running tap water for 5 min before sections were immersed in Eosin (01650, HistoLab, Sweden) for 30 seconds. Sections were dehydrated and mounted in PERTEX (00811, HistoLab, Sweden).

### Gene expression profiling by microarray

Cerebellar tissue, cleared from meninges, from mutant embryos (*Pdgfc^-/-^; Pdgfra^GFP/+^*) were compared to all other embryos in the same litter (*Pdgfc^+/-^; Pdgfra^GFP/+^, Pdgfc^+/+^; Pdgfra^GFP/+^*, *Pdgfc^-/-^, Pdgfc^+/-^* and *Pdgfc^+/+^*). Three litters were included; at E13.5 (3 mutants and 5 controls), at E14.5 (2 mutants and 7 controls, and at P0 (4 mutants and 9 controls). From the P0-litter, the meninges were isolated and analysed separately. Dissected tissue was stored in RNAlater®(Ambion) before RNA isolation using the RNeasy microkit (Qiagen). RNA quality was checked in a 2100 BioAnalyzer (AgilentTechnologies, Santa Clara, CA, USA). Transcription profiling was performed with the Gene Chip Mouse Gene 1.0ST array, at the Bioinformatics and Expression analysis core facility, Karolinska Institute, Sweden (http://apt.bea.ki.se). Affymetrix raw data was normalized using PLIER algorithms (AffymetrixTechnical Note, Guide to Probe Logarithmic Intensity Error Estimation, http://affymetrix.com/support/technical/technotesmain.affx). Cut-off values were >2 in fold change with a P-value <0.05 for the t-test.

The entire microarray data set has been deposited in the NCBI GeneExpression Omnibus database (www.ncbi.nlm.nih.gov/geo/, accession number GSE77049 and GSE202835).

## Acknowledgements

This project was financed by Svenska Läkaresällskapet (2006-18466), Åke Wibergs Stiftelse (362565719, 946216308), Stiftelsen Lars Hiertas Minne, MagnBergvalls Stiftelse, Karolinska Institutet, Swedish Research Council (C.B.: 2015-00550), the European Research Council (C.B.: AdG294556), the Leducq Foundation (C.B.: 14CVD02), Swedish Cancer Society (C.B.:150735), Knut and Alice Wallenberg Foundation (C.B.: 2015.0030).

We acknowledge Jana Chmielniakova, Pia Petersson and Cecilia Olsson for technical assistance with animals.

## Supplementary images

**Fig. S1.**
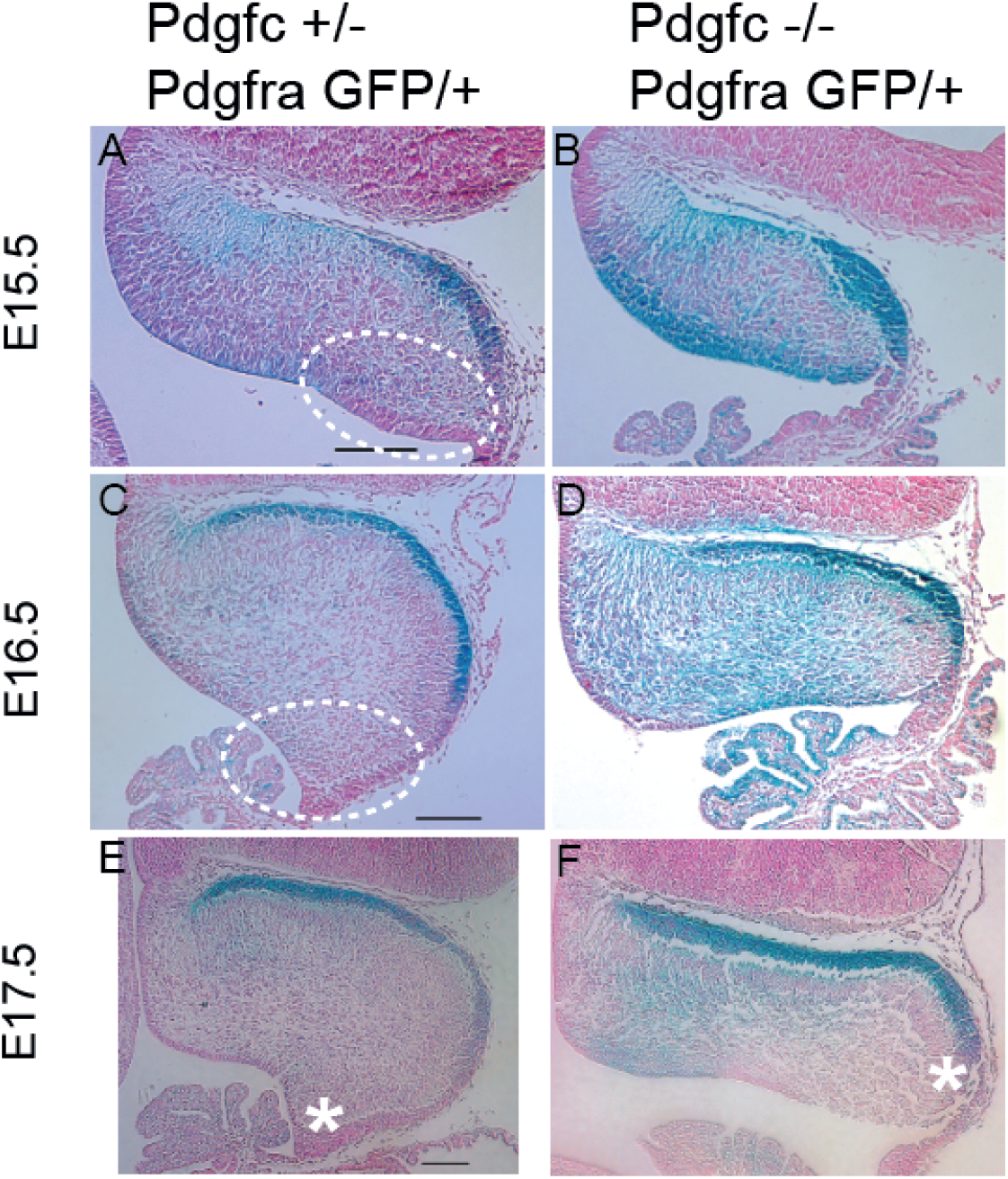
Lack of expansion in the rhombic lip area in *Pdgfc^-/-^; Pdgfra^GFP/+^* mice. X-gal staining as a reporter for *Pdgfc* expression in mid-sagittal sections of cerebellum of prenatal pups. (A, C, E) In control mice, the area close to the rhombic lip expanded and the cerebellum acquired a rounded shape. *Pdgfc* expression was restricted to granule cells in the EGL. (B, D, F) Cerebellum in mutant mice remained flat and elongated. As a result, the rhombic lip and the choroid plexus was dorsally located (asterisk). Each section is a representative of >3 mice.

**Fig. S2.**
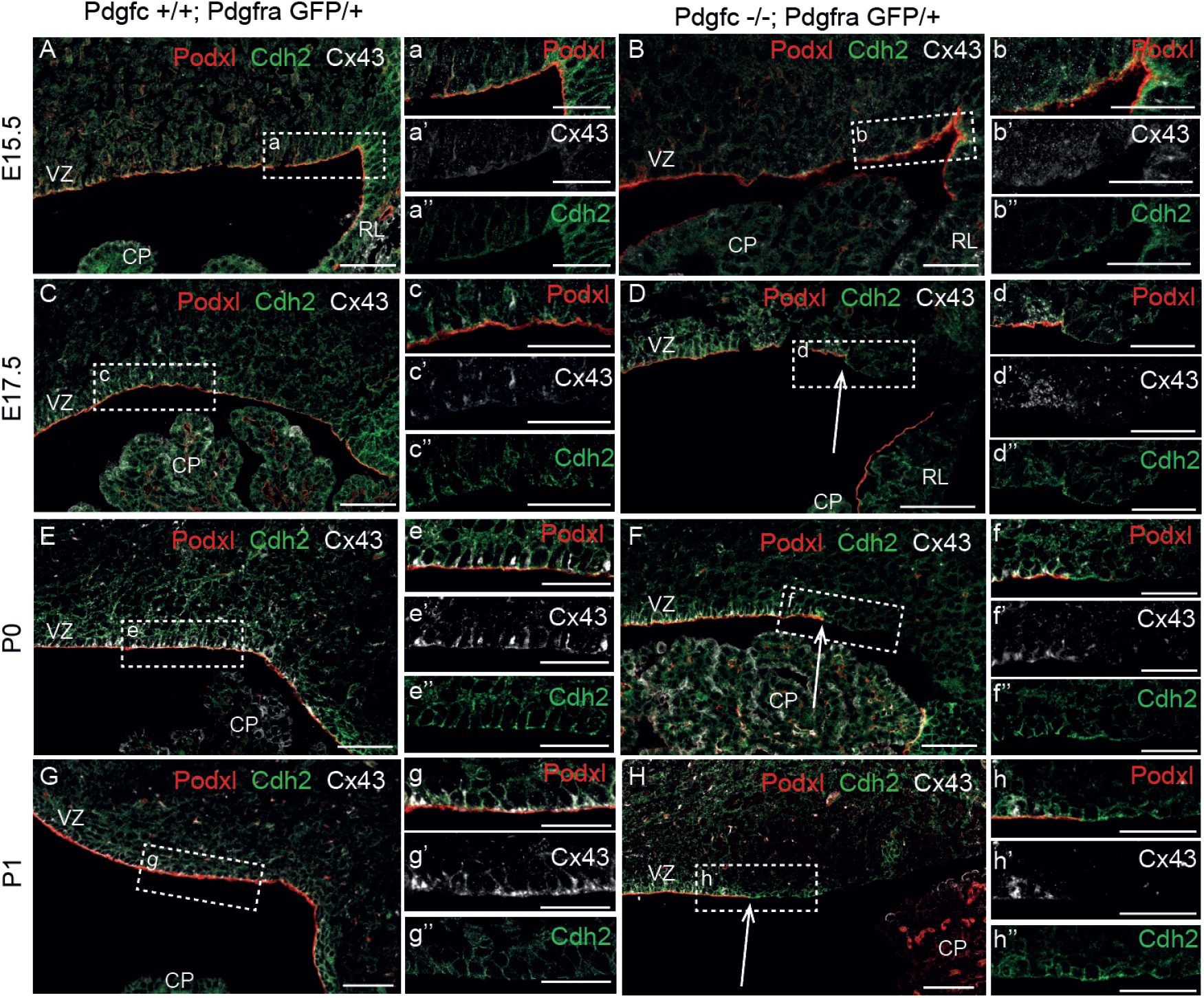
Alterations in expression of ependyma-specific markers. Immunofluorescent staining of Cdh2 and Cx43 in mid-sagittal sections of the ventricular zone at E15.5, E17.5, P0 and P1. (A, C, E, G) In control mice, podocalyxin was expressed in the apical side of ependymal cells. Cdh2 and Cx43 expression was expressed paracellular of the ependymal cells. (a, c, e, g) High magnification view. (D, F, H) From E17.5, podocalyxin expression did not cover the whole ventricular zone. Expression of Cx43 was lost and expression of Cdh2 was reduced. VZ-ventricular zone, CP-choroid plexus, RL-rhombic lip. Arrows indicate where podocalyxin expression was lost. Scalebar (A-H) 50 *μ*m, (a-h) 30 *μ*m. Each section is a representative of >3 mice.

## Supplementary Tables

**Table S1.**
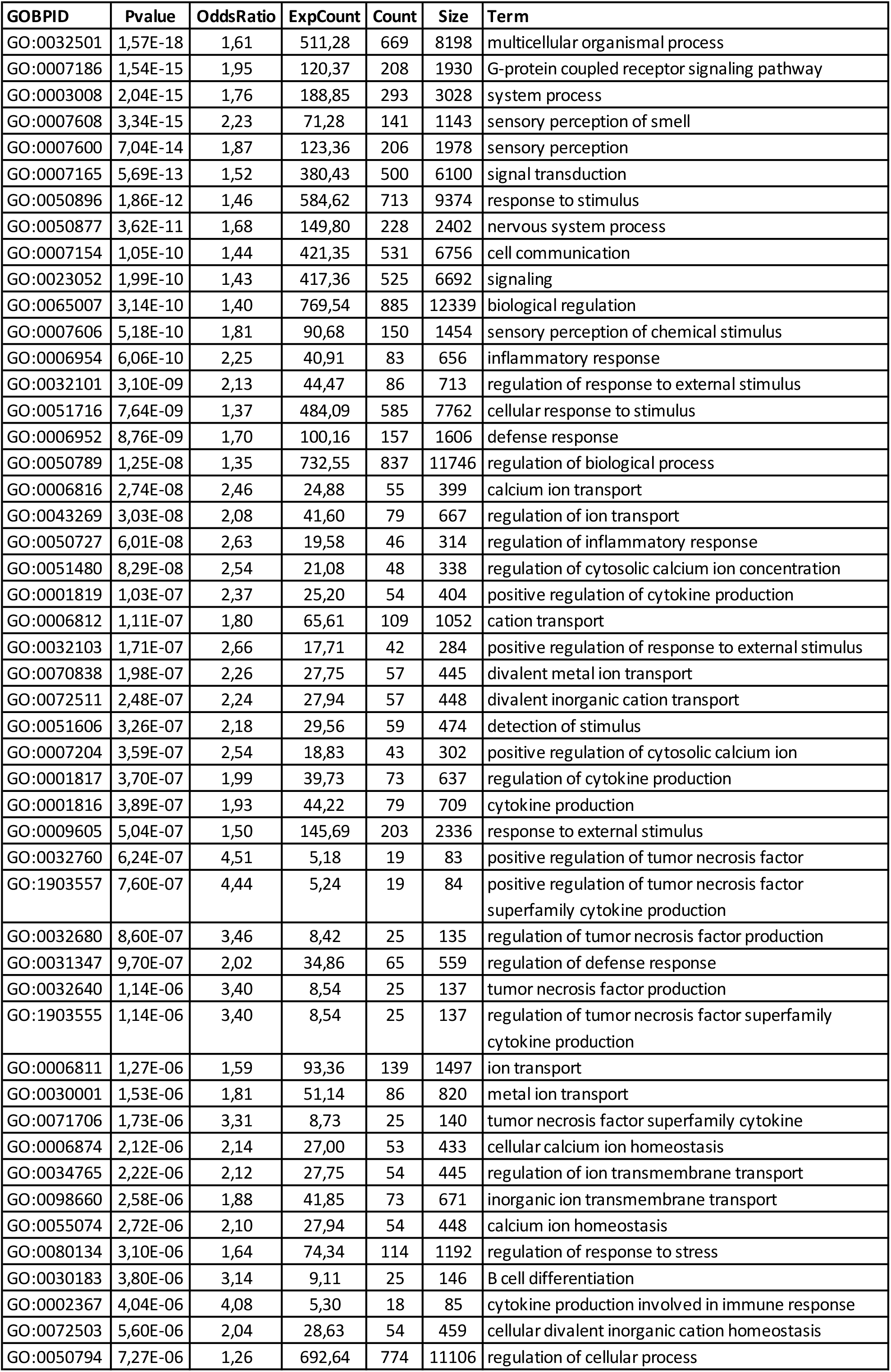
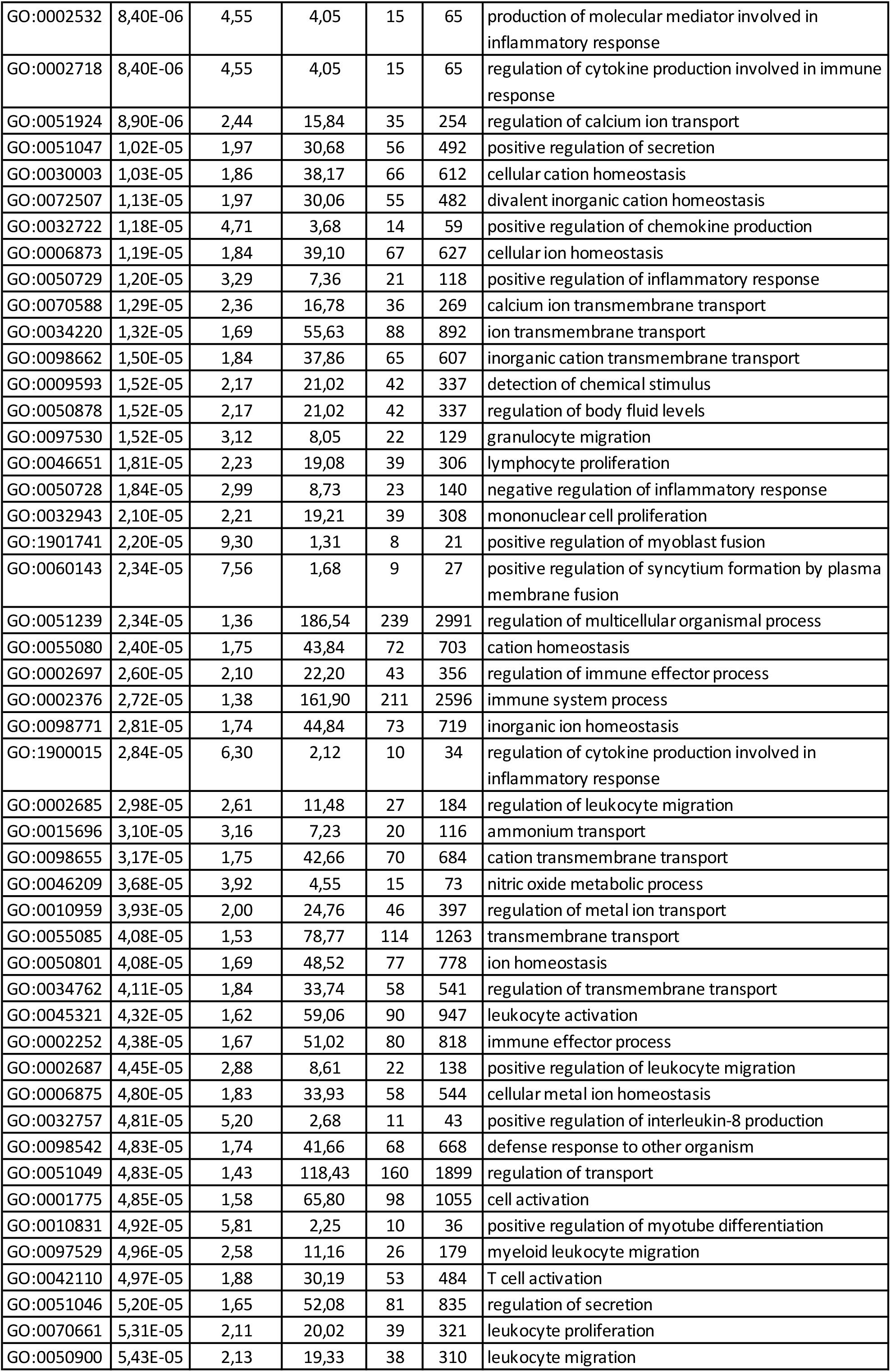
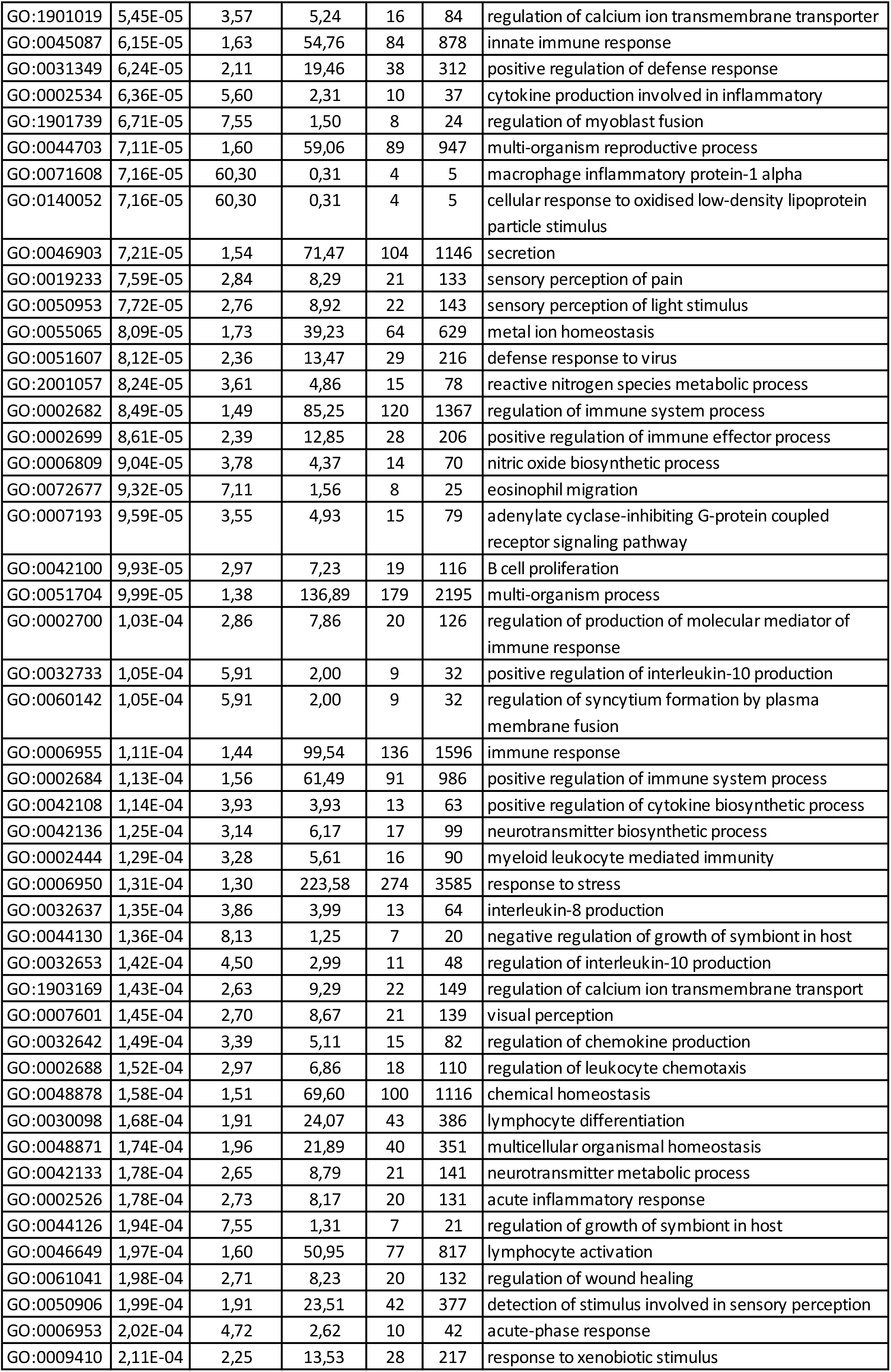
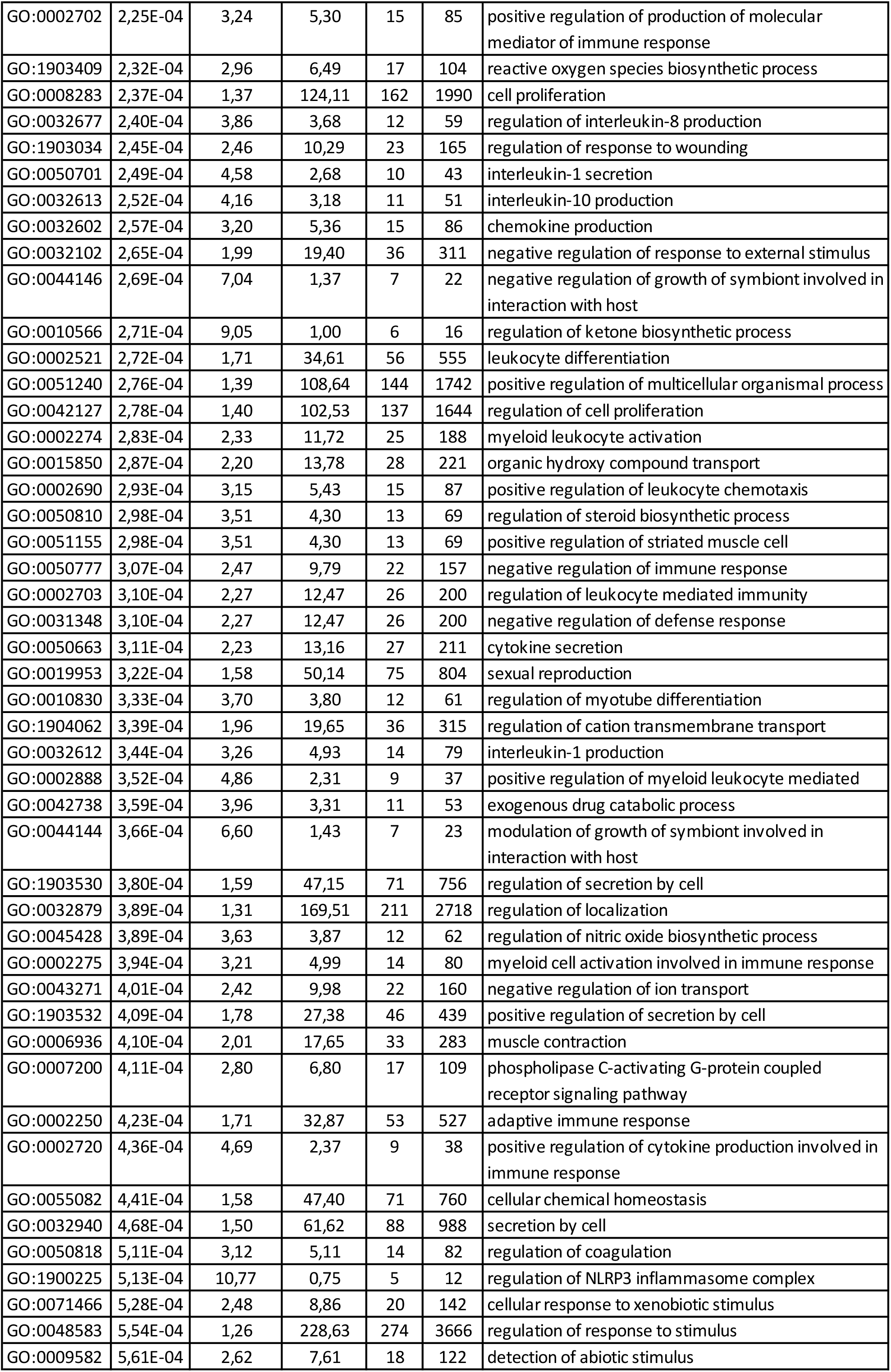
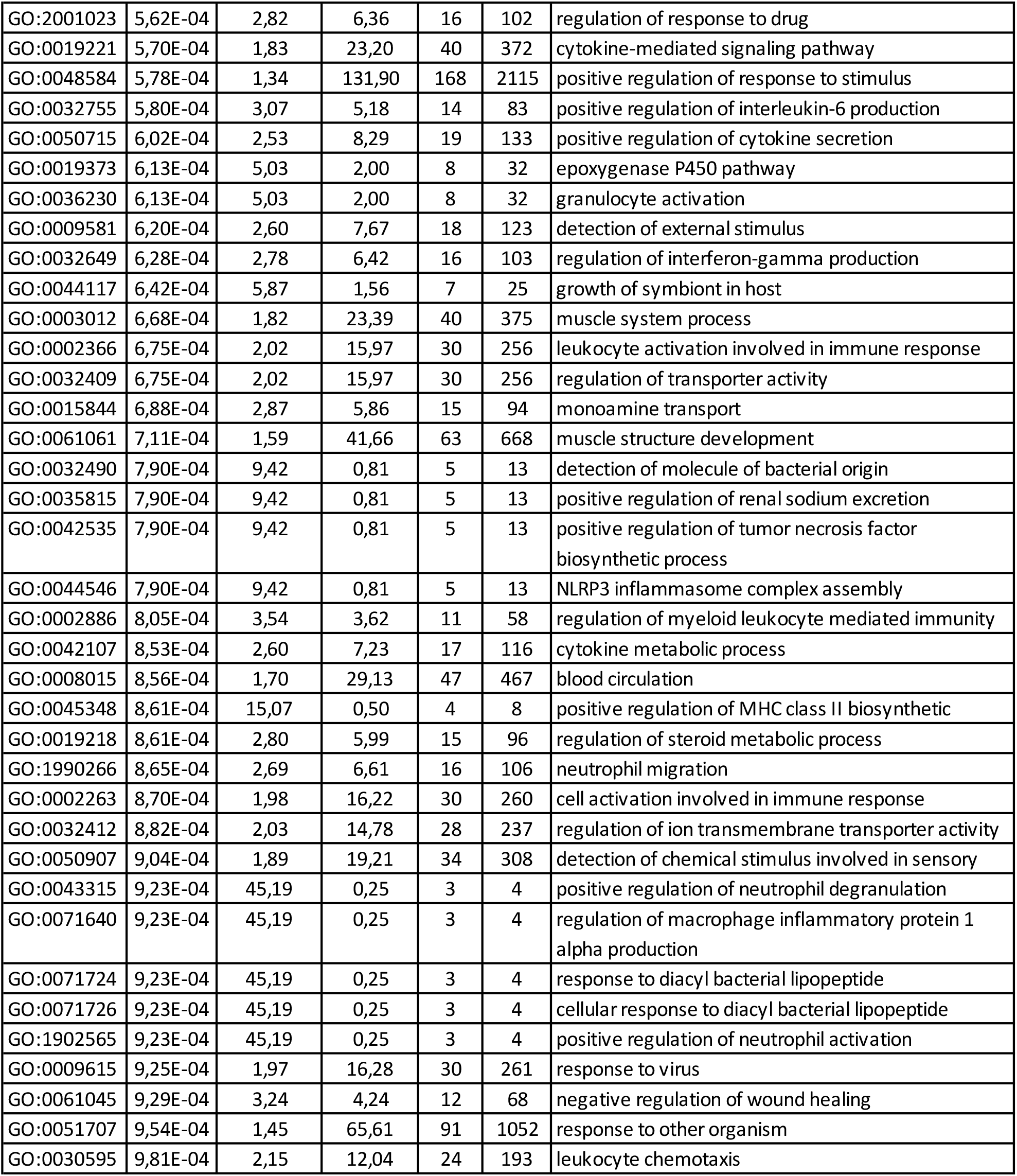
GO terms for downregulated differentially expressed genes in the cerebellum of E14.5 *Pdgfc^-/-^; Pdgfra^GFP/+^* mice. Based on a litter with two mutant and seven control pups. Cut off fold change >2 and p<0.05.

**Table S2.**
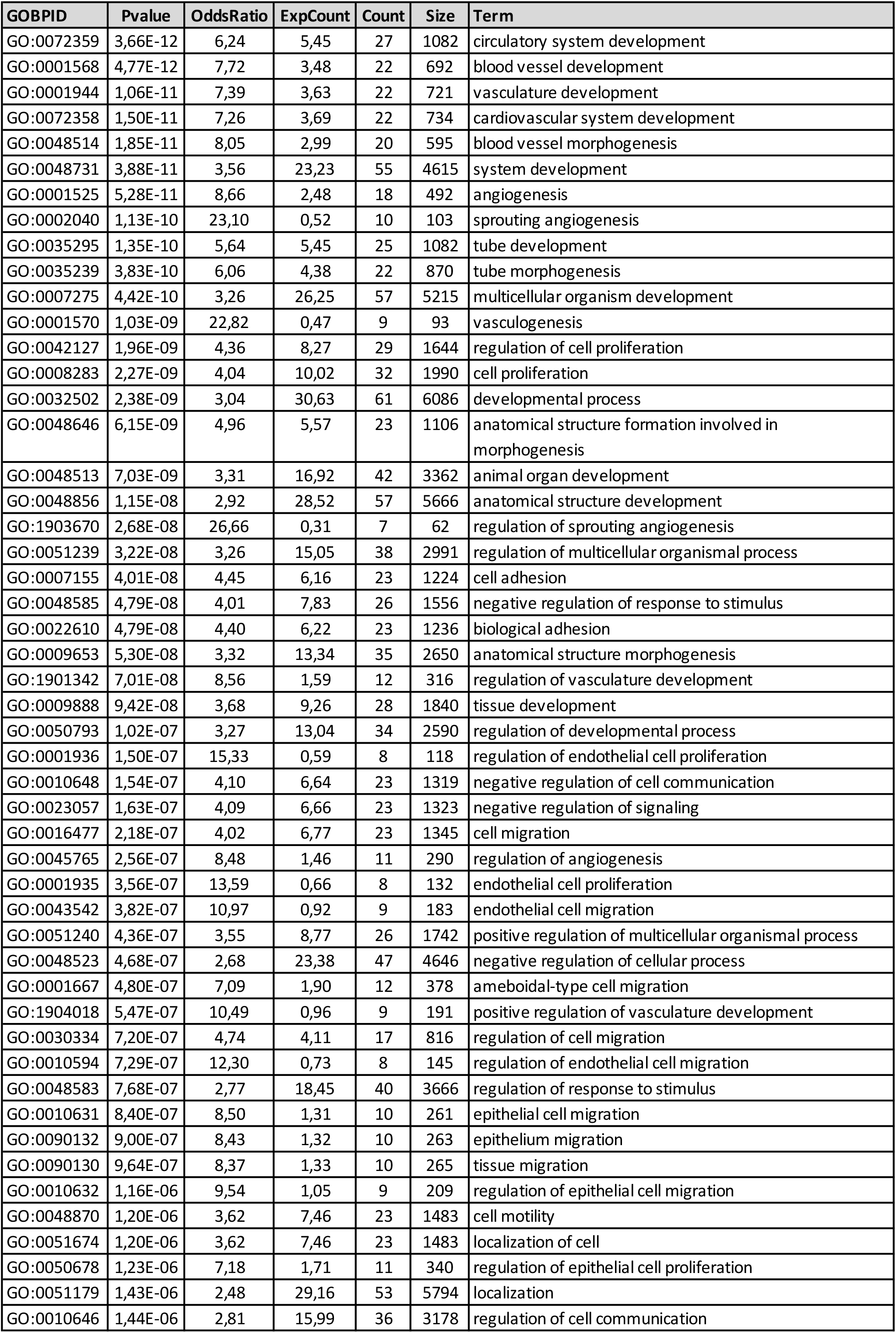
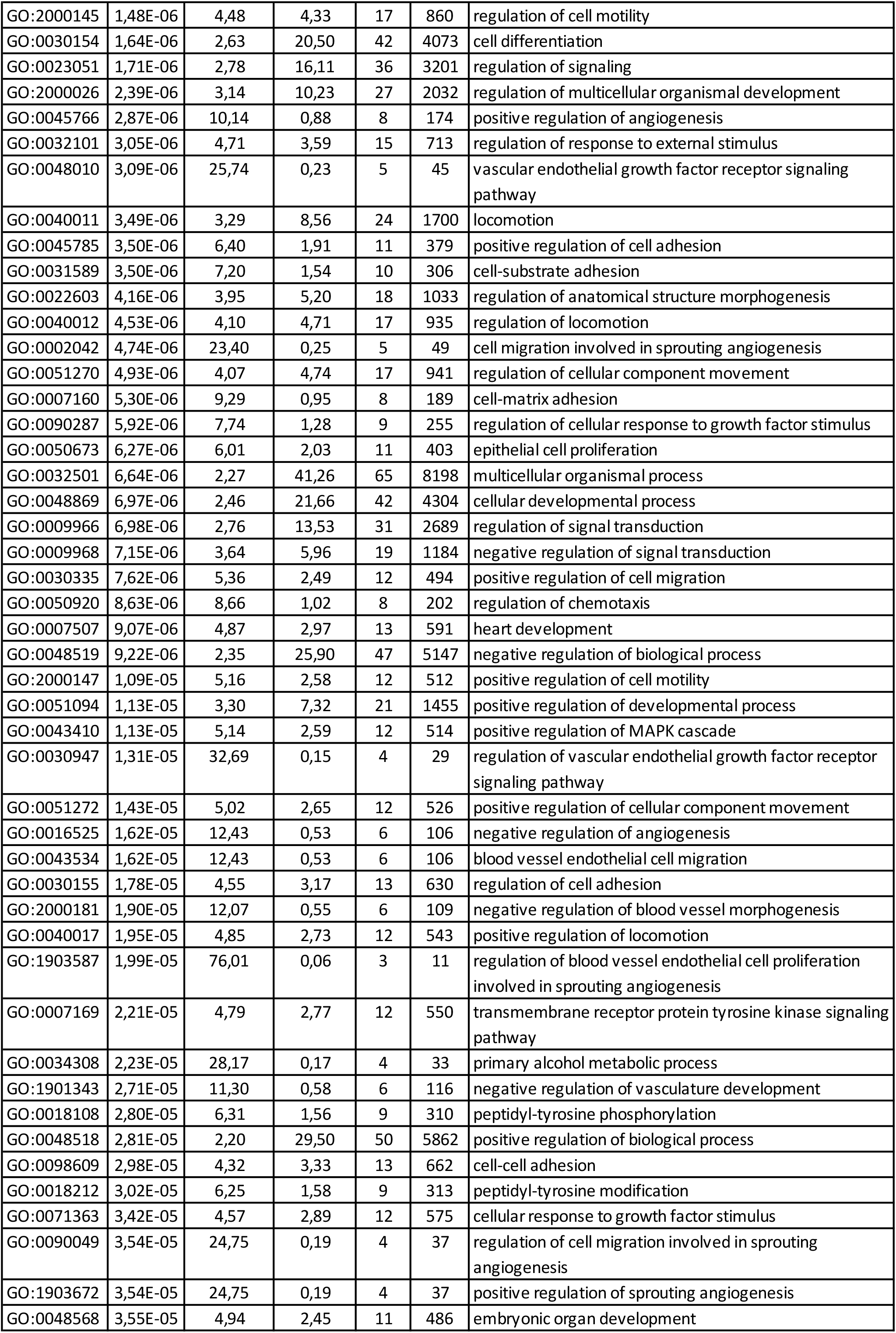
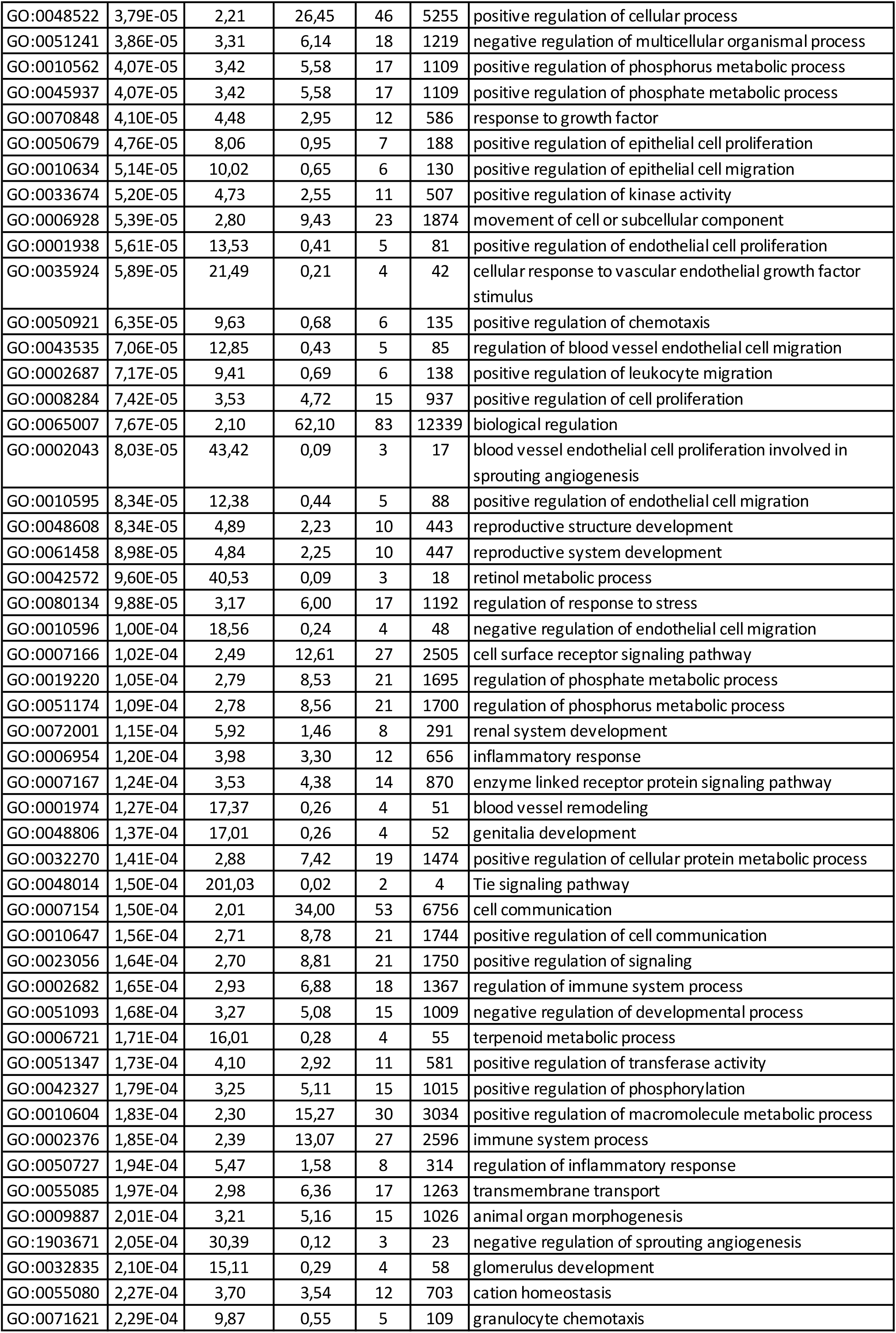
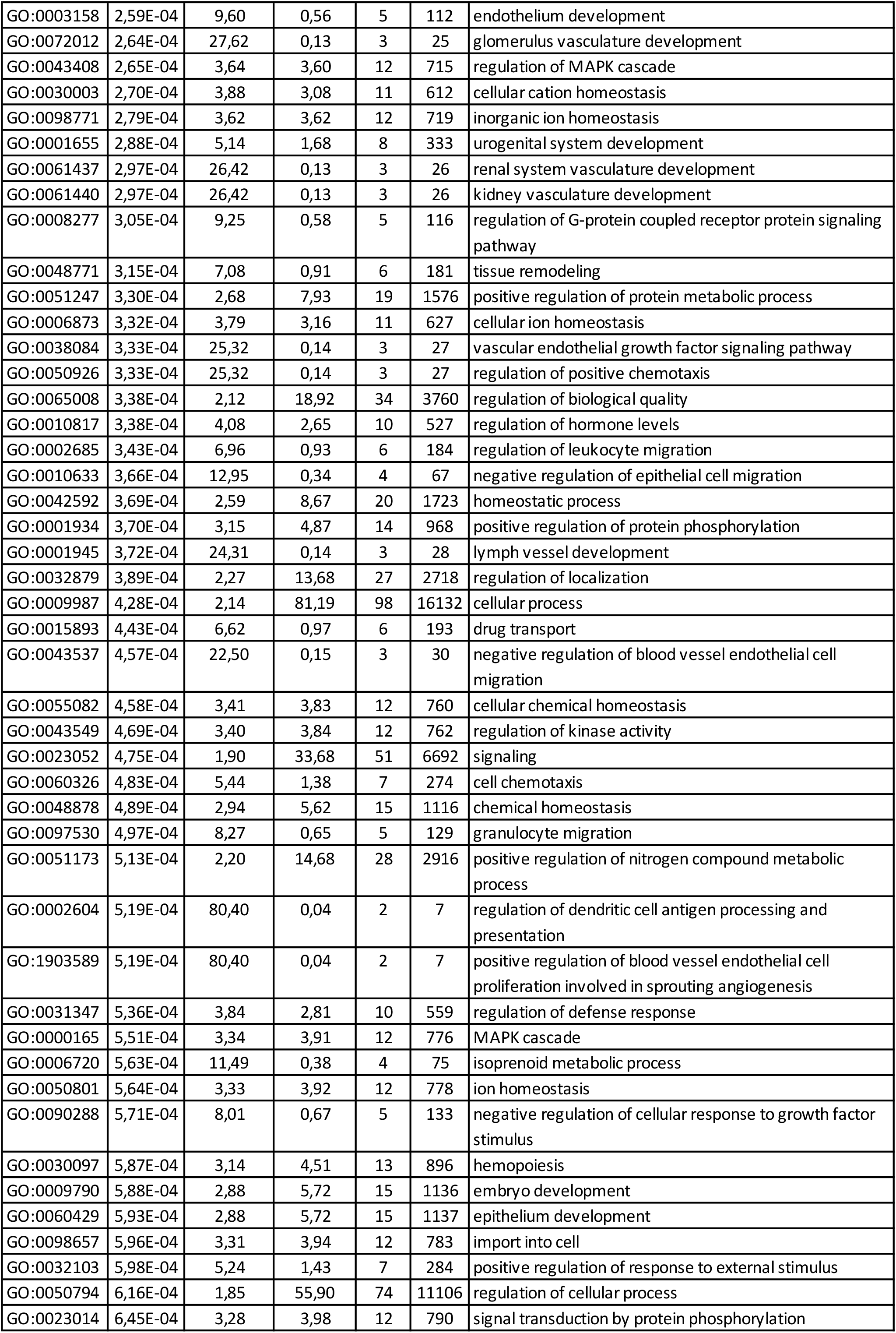
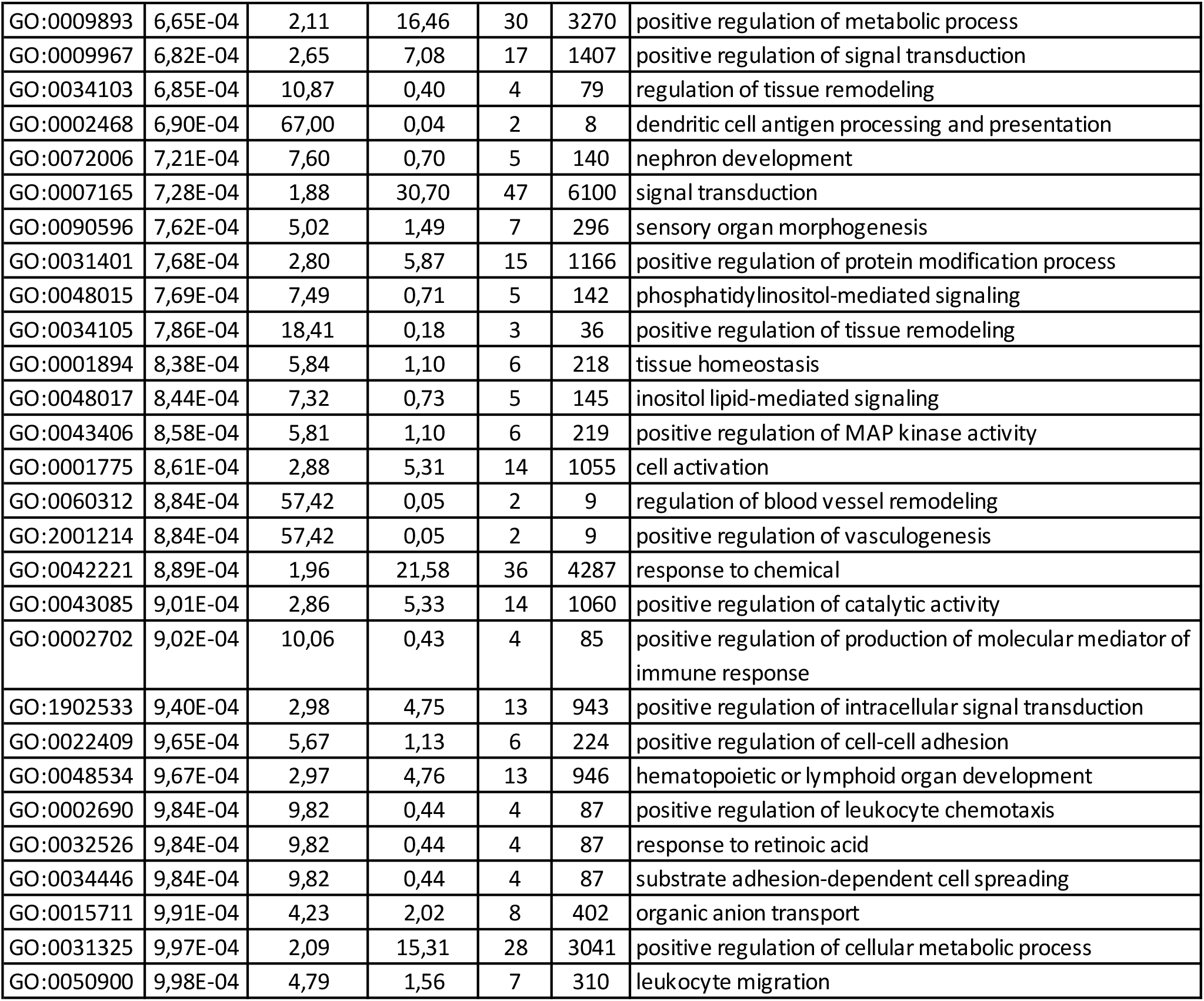
GO terms for downregulated differentially expressed genes in cerebellar meninges from P0 *Pdgfc^-/-^; Pdgfra^GFP/+^* mice. Based on a litter with four mutant and nine control pups. Cut off fold change >2 and p<0.05.

**Supplementary table 3:**
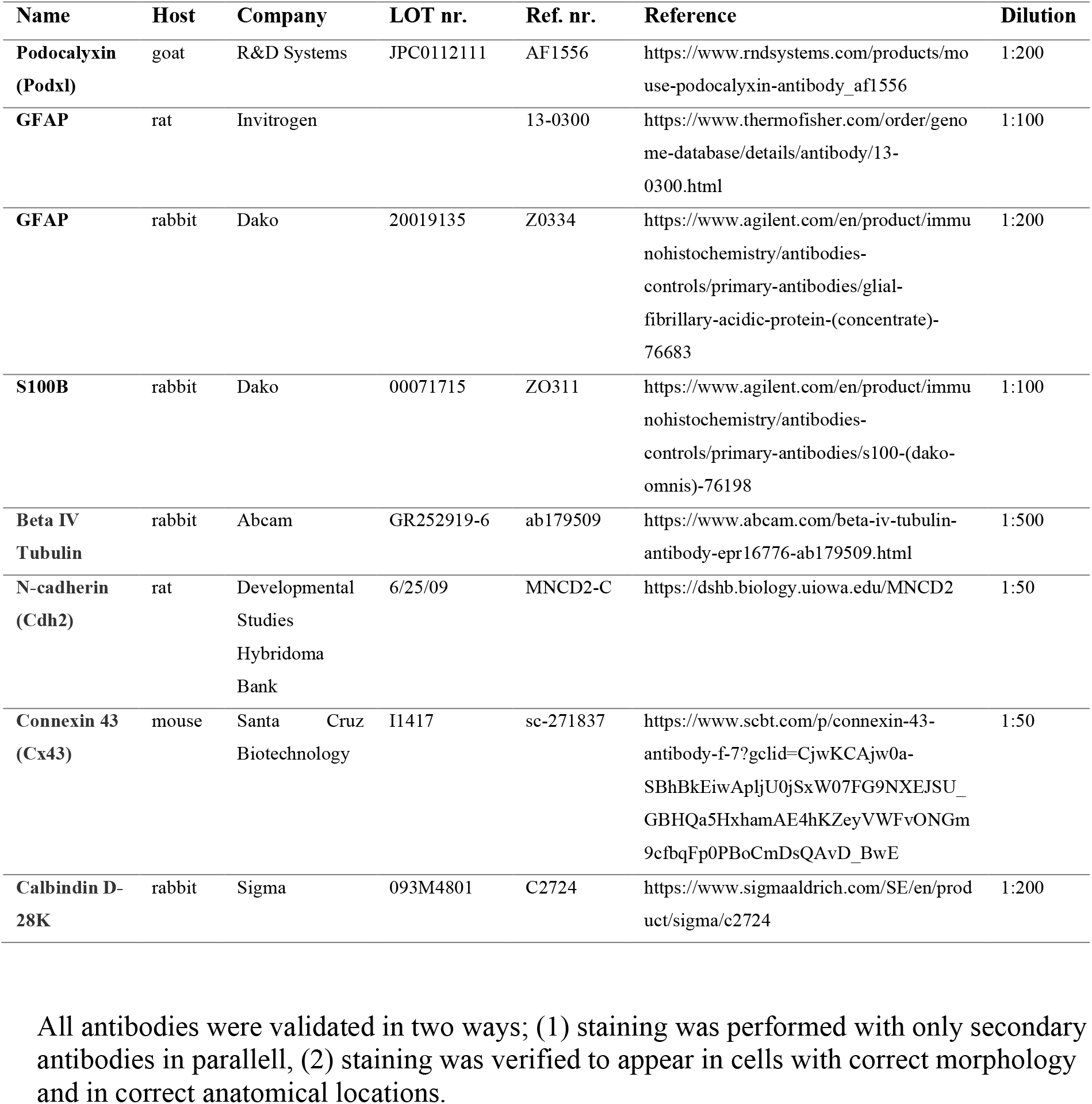
Antibodies and dilutions. Primary antibodies

